# AQ-seq: Accurate quantification of microRNAs and their variants

**DOI:** 10.1101/339606

**Authors:** Haedong Kim, Jimi Kim, Kijun Kim, Hyeshik Chang, Kwontae You, V. Narry Kim

## Abstract

MicroRNAs (miRNAs) modulate diverse biological and pathological processes via post-transcriptional gene silencing. High-throughput small RNA sequencing (sRNA-seq) has been widely adopted to investigate the functions and regulatory mechanisms of miRNAs. However, accurate quantification has been limited owing to the severe ligation bias in conventional sRNA-seq methods. Here we present a high-throughput protocol, termed AQ-seq (accurate quantification by sequencing), that utilizes adapters with terminal degenerate sequences and a high concentration of polyethylene glycol (PEG), which removes the ligation bias during library preparation. By accurately measuring miRNAs and their variants (known as isomiRs), we identify alternatively processed miRNAs and correct the 5′ end usage and strand preference that have been misannotated. We also uncover highly modified miRNAs that are uridylated and adenylated. Taken together, AQ-seq reveals the complexity of the miRNA isoform landscape, allowing us to refine miRNA annotation and to advance our understanding of miRNA regulation. Furthermore, AQ-seq can be adopted to improve other ligation-based sequencing methods including crosslinking-immunoprecipitation-sequencing (CLIP-seq) and ribosome profiling (Ribo-seq).

## Introduction

MicroRNAs (miRNAs) are ~22 nt-long small non-coding RNAs that regulate gene expression by inducing deadenylation and translational repression of target mRNAs (Bartel 2009). Biogenesis of miRNA involves multiple steps. Primary miRNAs (pri-miRNAs) are synthesized by RNA polymerase II and subsequently cleaved by a nuclear ribonuclease (RNase) III enzyme Drosha, releasing small hairpin-shaped precursor miRNAs (pre-miRNAs) (Lee et al. 2003; Denli et al. 2004; Gregory et al. 2004; Han et al. 2004). Pre-miRNAs are exported to the cytoplasm (Yi et al. 2003; Lund et al. 2004), where they are further processed by a cytoplasmic RNase III enzyme Dicer into ~22 nt-long duplex (Bernstein et al. 2001; Grishok et al. 2001; Hutvagner et al. 2001; Knight and Bass 2001). The miRNA duplex is loaded onto an Argonaute (AGO) protein, out of which one strand (“guide”) remains as the mature miRNA while the other (“passenger”) gets expelled. The strand with uridine or adenosine at its 5′ end binds readily to the 5′ pocket in the Mid domain of AGO and is subsequently selected as the guide strand. The strand with thermodynamically unstable binding at its 5′ end, due to mismatches to the opposite strand, A-U pair, or G-U pair, is also easily inserted into the 5′ pocket of AGO (Khvorova et al. 2003; Schwarz et al. 2003; Frank et al. 2010; Suzuki et al. 2015). The guide strand base-pairs with the target mRNA mainly through the nucleotides 2-7 relative to the 5′ end of miRNA (so called “seed” sequence), and induces gene silencing in a sequence-specific manner (Bartel 2009).

During maturation, multiple miRNA isoforms (isomiRs) can be generated from a single pri-miRNA hairpin. Alternative cleavage by Drosha or Dicer produces isomiRs with different 5′ and/or 3′ ends (Wu et al. 2009; Chiang et al. 2010; Kim et al. 2017). The 5′ end variation is of particular importance because it changes the seed sequence and, hence, target specificity. Altered cleavage also changes the 5′ end nucleotide and stability of miRNA duplex, affecting strand selection. Therefore, the 5′ end variation can substantially influence target repertoires.

Another major source of isomiRs is the 3′ end modifications by terminal nucleotidyl transferases. The non-templated nucleotidyl addition (or “RNA tailing”) can occur at both the pre- and mature miRNA stages. RNA tailing modulates downstream processing and stability of miRNAs (Heo et al. 2008; Hagan et al. 2009; Heo et al. 2009; Heo et al. 2012; Lee et al. 2014; Thornton et al. 2014; Kim et al. 2015). For example, when mono-uridylation occurs on pre-let-7 with a 1-nt 3′ overhang (classified into “group II”), the U-tail extends the 3′ overhang to make an optimal substrate for Dicer, upregulating pre-let-7 processing (Heo et al. 2012). In contrast, oligo-uridylation of pre-let-7, which is induced by Lin28, blocks pre-let-7 processing and induces its degradation (Heo et al. 2008; Hagan et al. 2009; Heo et al. 2009).

High-throughput small RNA sequencing (sRNA-seq) has been widely adopted to discover and quantify functionally important miRNAs and their variants. sRNA-seq can profile miRNAs at a single nucleotide resolution and detect isomiRs without prior knowledge. However, accurate miRNA profiling has been difficult because certain miRNAs are favored over others in an enzyme-dependent ligation reaction due to the preference of RNA ligases for some sequences and structures, leading to a skewed representation of miRNAs. This can severely compromise quantitative analysis of end heterogeneity and strand preference of a given miRNA (Hafner et al. 2011; Jayaprakash et al. 2011; Sun et al. 2011; Sorefan et al. 2012; Zhuang et al. 2012; Zhang et al. 2013; Song et al. 2014; Dard-Dascot et al. 2018). Recent studies have sought to minimize the ligation bias by adopting randomized adapters, presuming that the increased diversity of adapter sequences raises chances of capturing miRNAs with various sequences (Jayaprakash et al. 2011; Sun et al. 2011; Sorefan et al. 2012; Zhuang et al. 2012; Zhang et al. 2013; Fuchs et al. 2015; Xu et al. 2015; Dard-Dascot et al. 2018). Another approach was to apply polyethylene glycol (PEG), which facilitates ligation reaction via molecular crowding effect (Harrison and Zimmerman 1984; Zhang et al. 2013; Song et al. 2014; Xu et al. 2015; Shore et al. 2016; Dard-Dascot et al. 2018). Each strategy has shown some degree of success in ameliorating the ligation bias, yet the optimal condition for their combinatorial use has not been investigated.

In this study, we systematically evaluate the sequencing bias pervasive in sRNA-seq, and develop a bias-minimized protocol to perform a comprehensive study on miRNA heterogeneity.

The results correct previously misannotated miRNAs, and reveal novel maturation events, including alternative processing, uridylation, and strand selection.

## Results

### Small RNA sequencing optimization using spike-ins

To assess the extent of bias in sRNA-seq data, we developed spike-in controls which consist of 30 synthetic RNAs of 21-23 nt (Supplemental Table S1). The 30 exogenous RNAs were added to total RNA at equimolar concentrations, and subjected to sRNA-seq library preparation (Fig. 1A). We generated a sequencing library using the conventional method, called TruSeq, and evaluated bias by calculating relative amounts of spike-ins. We observed a strikingly skewed representation of spike-ins in the sequencing result, which reflects severe bias (Fig. 1B, top left panel). In light of this observation, we set out to optimize the sRNA-seq protocol by introducing randomized adapters containing four degenerate nucleotides. The randomized adapters reduced the bias but not to a satisfying degree (Fig. 1B, top right panel). A high concentration of PEG (20%) also ameliorated the ligation bias (Fig. 1B, bottom left panel), but did not fully resolve the problem. Some spike-ins were still grossly overestimated; ~6 out of 30 spike-ins still accounted for more than 50% of total spike-in reads when only randomized adapters or only PEG was applied. Thus, we tested various combinations of randomized adapters and PEG (Supplemental Fig. S1, lanes 2, 4-15), and found that either applying PEG only to the 3′ adaptor ligation step or adopting less than 20% of PEG - as being used in currently available protocols (Zhang et al. 2013; Xu et al. 2015; Dard-Dascot et al. 2018) - is not sufficient to abolish the ligation bias. Substantial improvement was achieved when we included 20% PEG at both 3′ and 5′ adapter ligation reactions, in combination with randomized adapters (Fig. 1A). Hereafter we refer to this protocol as AQ-seq (accurate quantification by sequencing).

**Figure 1.**
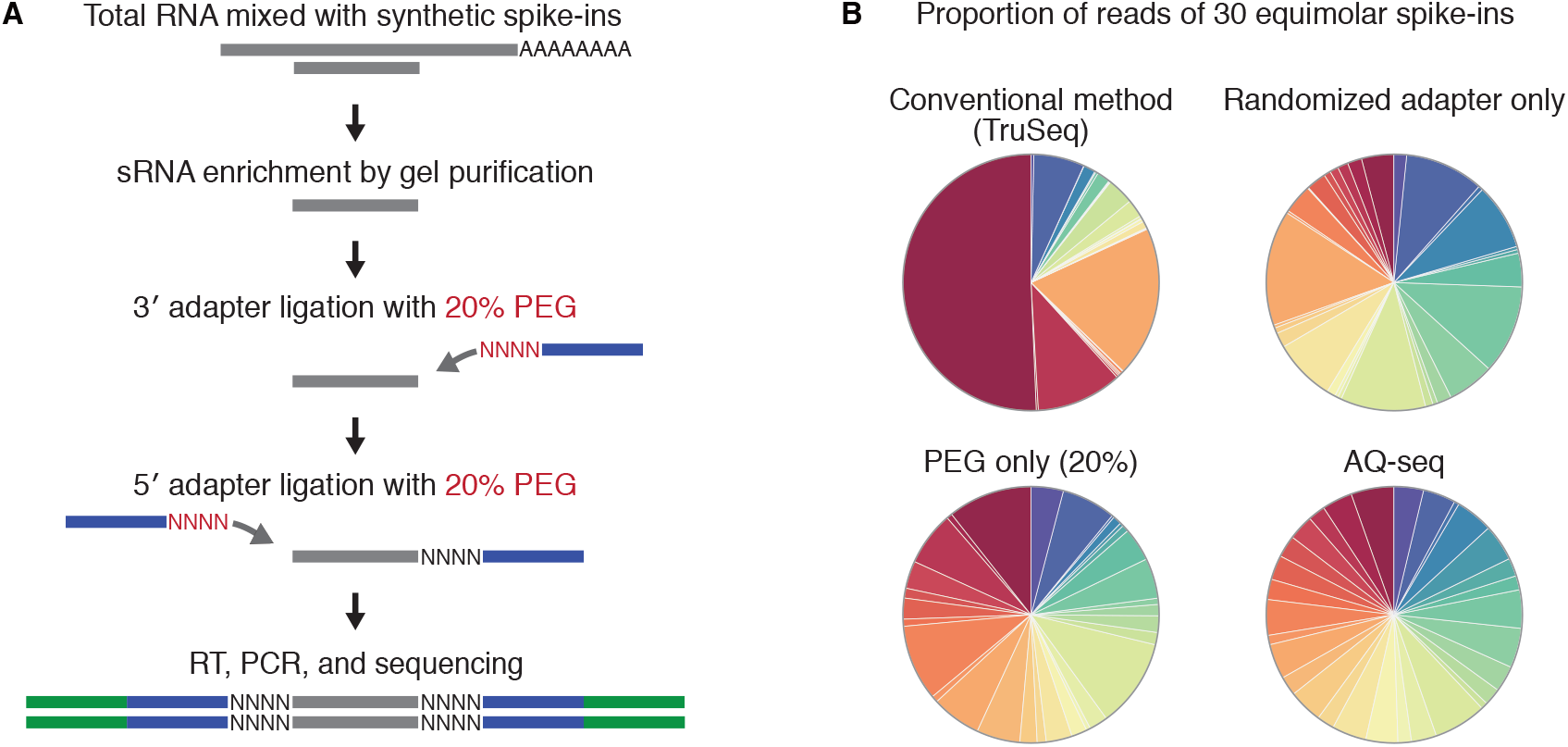
AQ-seq substantially reduces sequencing bias. (A) Schematic outline of AQ-seq library preparation method. (B) Proportion of 30 spike-ins detected by the indicated protocols.

### miRNA and isomiR profiles revealed by AQ-seq

To evaluate the results from AQ-seq, we examined miRNA abundance in two human cell lines, HEK293T and HeLa, determined by AQ-seq in comparison with that by TruSeq. While miRNA and isomiR profiles from replicates within each method are highly reproducible (Supplemental Fig. S2), the profiles produced by the two methods correlated poorly with each other (Fig. 2). Notably, AQ-seq detected ~1.5 fold more miRNAs than TruSeq did, indicating that AQ-seq is more sensitive than TruSeq is (Fig. 2A, B).

**Figure 2.**
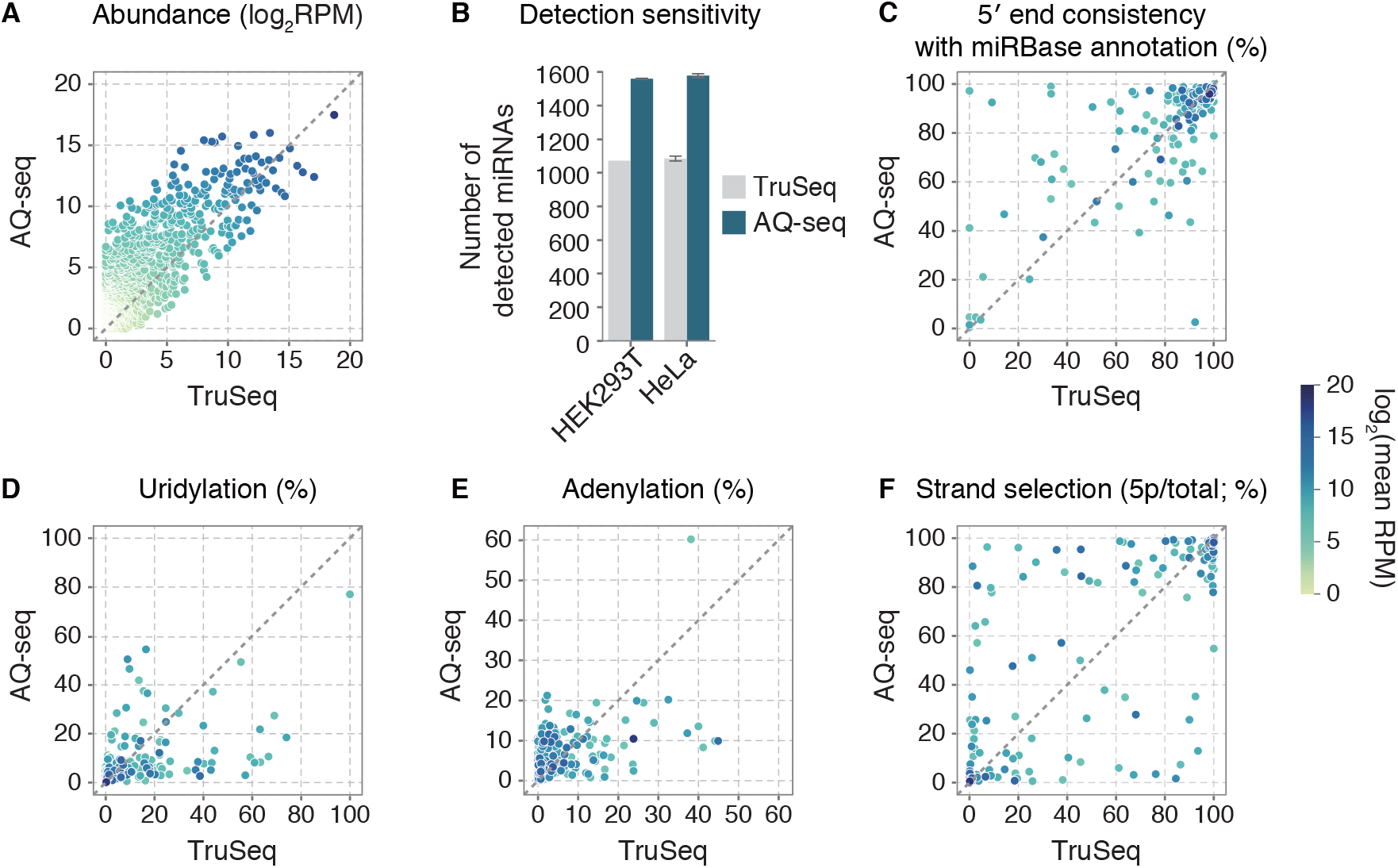
There are large differences in miRNA and isomiR profiles between AQ-seq and TruSeq. (A-F) Comparison of miRNA and isomiR profiles between AQ-seq and TruSeq in HEK293T cells. Abundant miRNAs (>100 RPM in AQ-seq or TruSeq) were included in each analysis, unless otherwise indicated. RPM, reads per million. (A) Expression profiles. No abundance filter was applied. (B) The number of detected miRNAs in the indicated methods. No abundance filter was applied. Bars indicate mean +/− standard deviations (s.d.) (n=2). (C) The portion of reads whose 5′ ends start at the same position with that of miRBase for a given miRNA (Kozomara and Griffiths-Jones 2014). (D and E) Terminal modification frequencies by uridylation (D) or adenylation (E). All terminally modified reads were counted regardless of the tail length. (F) Relative portion of 5p and 3p strands calculated by [5p]/([5p]+[3p]), where brackets mean read counts.

We next examined the isomiR profiles. The 5’-isomiR profile from AQ-seq was markedly different from that of TruSeq (Fig. 2C). As for the 3′ terminal modification, highly uridylated miRNAs were identified by both methods, but they were inconsistent (Fig. 2D). Adenylation frequencies were mostly estimated to be lower than those determined by TruSeq (Fig. 2E). Lastly, the strand ratios of numerous miRNAs (5p vs. 3p) differ greatly between the two methods (Fig. 2F). Collectively, AQ-seq provides the miRNA profiles drastically different from those obtained by TruSeq.

### AQ-seq detects the 5′ ends that are previously misannotated or undefined

The 5′ end of the guide strand is particularly important for miRNA functionality because the “seed” sequence located at 2-7 nt position relative to the 5′ end of miRNA dictates the specificity of target recognition (Bartel 2009). Since AQ-seq and TruSeq reported inconsistent 5′ ends for many miRNAs (Fig. 2C, 3A; 17 in HEK293T and 11 in HeLa cells among miRNAs with reads per million (RPM) over 100), we investigated which method is more reliable in detecting the true start site of miRNAs. For validation, we selected miR-222-5p due to the discrepancy between the two methods (Fig. 3B). TruSeq detected the 5′ end of miR-222-5p as annotated in the miRBase, while AQ-seq revealed a different 5′ end that is shifted by 2 nt. Of note, the end detected by AQ-seq matches the Drosha cleavage site recently identified by formaldehyde crosslinking, immunoprecipitation, and sequencing (fCLIP-seq) (Kim et al. 2017) (Fig. 3B, upper panel). To reveal the 5′ terminus of miR-222-5p, we performed primer extension experiment (Fig. 3B, lower panel). The result showed that the 5′ end of miR-222-5p was indeed correctly identified by AQ-seq and that the miRBase needs to be corrected.

**Figure 3.**
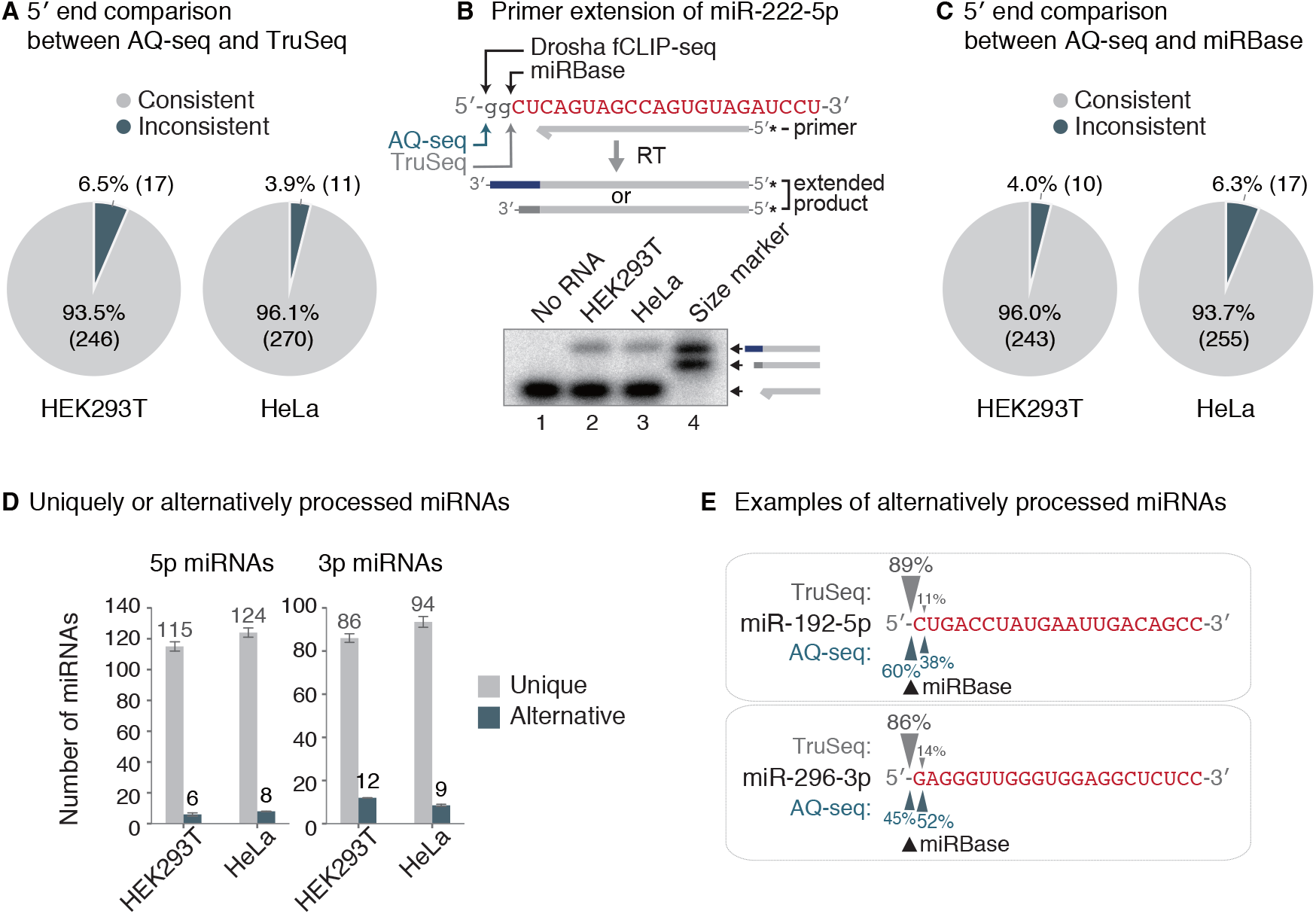
AQ-seq identifies misannotated 5′ ends and previously undetected alternative ends. (A) Comparison of miRNA 5′ ends between AQ-seq and TruSeq. The predominant end for a given miRNA among abundant miRNAs (>100 RPM in AQ-seq or TruSeq) was analyzed. (B) Primer extension of miR-222-5p. Identified 5′ ends of miR-222-5p from two sRNA-seq data (AQ-seq and TruSeq) and two resources (Drosha fCLIP-seq and miRBase) are indicated by arrows. The sequence of miR-222-5p deposited in miRBase is colored in red with upper cases. Synthesized oligonucleotides complementary to miR-222-5p with or without 2 nt-extended 5′ end were used as size references. Asterisks mark radiolabeled terminal phosphates. (C) Comparison of miRNA 5′ ends between AQ-seq and miRBase. Relatively abundant miRNAs (>100 RPM in AQ-seq) were included in this analysis. (D) The number of miRNAs with unique or multiple 5′ ends. Since the 5′ ends of 5p and 3p strands are processed by Drosha and Dicer, respectively, they were separately analyzed. If more than two 5′ ends of a given miRNA were detected and each of them account for >30% of total reads, the miRNA was considered as an alternatively processed miRNA. Otherwise, the miRNA was considered as a uniquely processed miRNA. (E) Illustration of 5′ end usage of two alternatively processed miRNAs, miR-192-5p and miR-296-3p. The portion of sequencing reads with the indicated 5′ end from HEK293T cells is denoted. The sequences of each miRNA deposited in miRBase are colored in red with upper cases.

This led us to examine the 5′ end discrepancy between miRBase and AQ-seq data. We found that about 4-6% of miRNAs detected in AQ-seq (RPM >100) have 5′ ends different from those annotated in miRBase (Fig. 3C). Comparison with the Drosha fCLIP-seq data indicates that AQ-seq detected the same termini as those from fCLIP-seq (Supplemental Fig. S3). Note that the 5′ end of 3p miRNA is determined by Dicer, and fCLIP data for Dicer is not currently available. Collectively, these results demonstrate that AQ-seq reliably captures the 5′ ends of miRNAs and offers an opportunity to correct previously misannotated 5′ ends.

Moreover, AQ-seq data allowed us to detect alternatively processed miRNAs (Fig. 3D). Although the majority of miRNAs appeared to be precisely cleaved at one position generating highly homogenous 5′ ends, certain miRNA hairpins produce substantial amounts of isomiRs with distinct 5′ ends. For instance, two isoforms of miR-192-5p and miR-296-3p are generated from alternative processing, which are rarely captured by TruSeq (Fig. 3E).

### AQ-seq identifies uridylated miRNAs

Intriguingly, a subset of miRNAs carry non-templated mono-uridine at high frequency (Fig. 4A). For instance, miR-551b-3p, miR-652-3p, miR-760, miR-30e-3p, and miR-324-3p are uridylated at more than ~50% frequency in HEK293T cells. Notably, uridylation frequencies measured by AQ-seq differ considerably from those by TruSeq (Supplemental Fig. S4). For experimental validation, we chose miR-652-3p which showed a large difference in its uridylation frequency and is abundant enough for Northern blot-based quantification (Fig. 4B). The major isoform of miR-652-3p was 1 nt longer than the reference sequence, suggesting that endogenous miR-652-3p is indeed highly mono-uridylated (Fig. 4B, left panel). Quantification of the band intensity confirmed that the longer isoform is more abundant than the shorter isoform (Fig. 4B, right panel).

**Figure 4.**
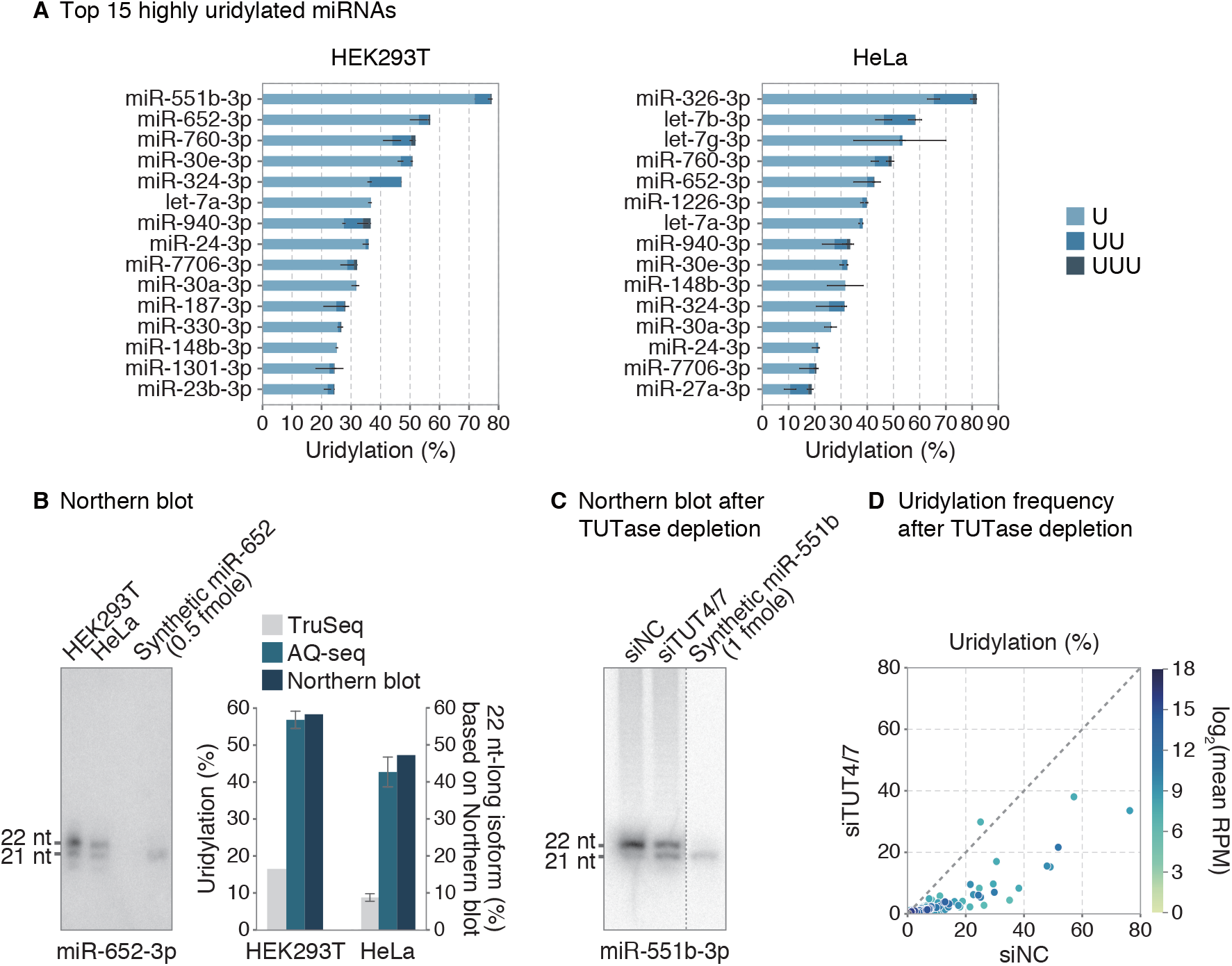
AQ-seq identifies highly uridylated miRNAs. (A) Top 15 highly uridylated miRNAs among abundant miRNAs (>100 RPM) in HEK293T and HeLa cells. The type of U-tail is indicated with different color. Light blue, blue, and navy refer to mono-uridylation (U), di-uridylation (UU), and tri-uridylation (UUU), respectively. Bars indicate mean +/− standard deviations (s.d.) (n=2). RPM, reads per million. (B) Northern blot of miR-652-3p detected in the indicated cell lines (left panel). Synthetic miR-652 duplex was loaded as a control. The proportion of the elongated isoform was quantified and compared to uridylation frequencies from TruSeq and AQ-seq (right panel). Bars indicate mean +/− standard deviations (s.d.) (n=2). (C) Northern blot of miR-551b-3p. The miRNA was detected 96 hr after transfection of indicated siRNAs in HEK293T cells. Synthetic miR-551b duplex was loaded as a control. (D) Uridylation frequencies calculated by AQ-seq after knockdown of TUT4 and TUT7 in HEK293T cells. Abundant miRNAs (>100 RPM) in the siNC-transfected sample were included in the analysis.

Next, we examined the enzyme(s) responsible for modification of the uridylated miRNAs. It has been reported that two terminal uridylyl transferases (TUTases), TUT4 (ZCCHC11), and TUT7 (ZCCHC6), uridylate pre-let-7 and some mature miRNAs (Hagan et al. 2009; Heo et al. 2009; Heo et al. 2012; Thornton et al. 2014; Kim et al. 2015). To test their involvement, we ectopically expressed pri-miR-551b after knockdown of TUT4 and TUT7. Northern blotting shows that, consistent with its high uridylation frequency (~80%) (Fig. 4A), the majority of miR-551b-3p was 1 nt longer than the synthetic miRNA mimic (21 nt). Depletion of TUT4/7 increased the short isoform, indicating that the 1-nt extension was due to mono-uridylation by TUT4/7 (Fig. 4C). We also performed AQ-seq after depletion of TUT4/7 and observed a global decrease in uridylation (Fig. 4D). These results indicate that the substrates of TUT4 and TUT7 are not limited to group II pre-miRNAs (pre-let-7, pre-miR-98, pre-miR-105-1, and pre-miR-449b) (Heo et al. 2012), recessed pre-miRNAs (Kim et al. 2015), and GUAG/UUGU-containing mature miRNAs (let-7, miR-10, miR-99/100, and miR-196 family) (Thornton et al. 2014). Of note, all of the highly uridylated miRNAs detected in our study are 3p miRNAs, originating from the 3′ strand of pre-miRNA (Fig. 4A). This confirms the previous notion that TUT4/7 act on pre-miRNAs prior to Dicer-mediated processing.

### AQ-seq data reveal strand preference

The most striking discrepancy between AQ-seq and TruSeq was found in strand preference (Fig. 2F). According to the AQ-seq data, miR-17, miR-106b, and miR-151a produce more 5p miRNAs than 3p miRNAs, but TruSeq gave the opposite results (Fig. 5A; Supplemental Fig. S5A). As for miR-423, 3p is more abundant than 5p according to AQ-seq whereas 5p is supposed to be dominant based on TruSeq and miRBase. Estimation of strand ratio by quantitative real-time PCR (RT-qPCR) was consistent with AQ-seq data rather than TruSeq data (Supplemental Fig. S5B). To further validate the strand ratio, we performed Northern blotting using synthetic miR-423 duplex of known quantity as a control. The strand ratio measured by Northern blot analysis matched well to those measured by AQ-seq and RT-qPCR (Fig. 5B).

**Figure 5.**
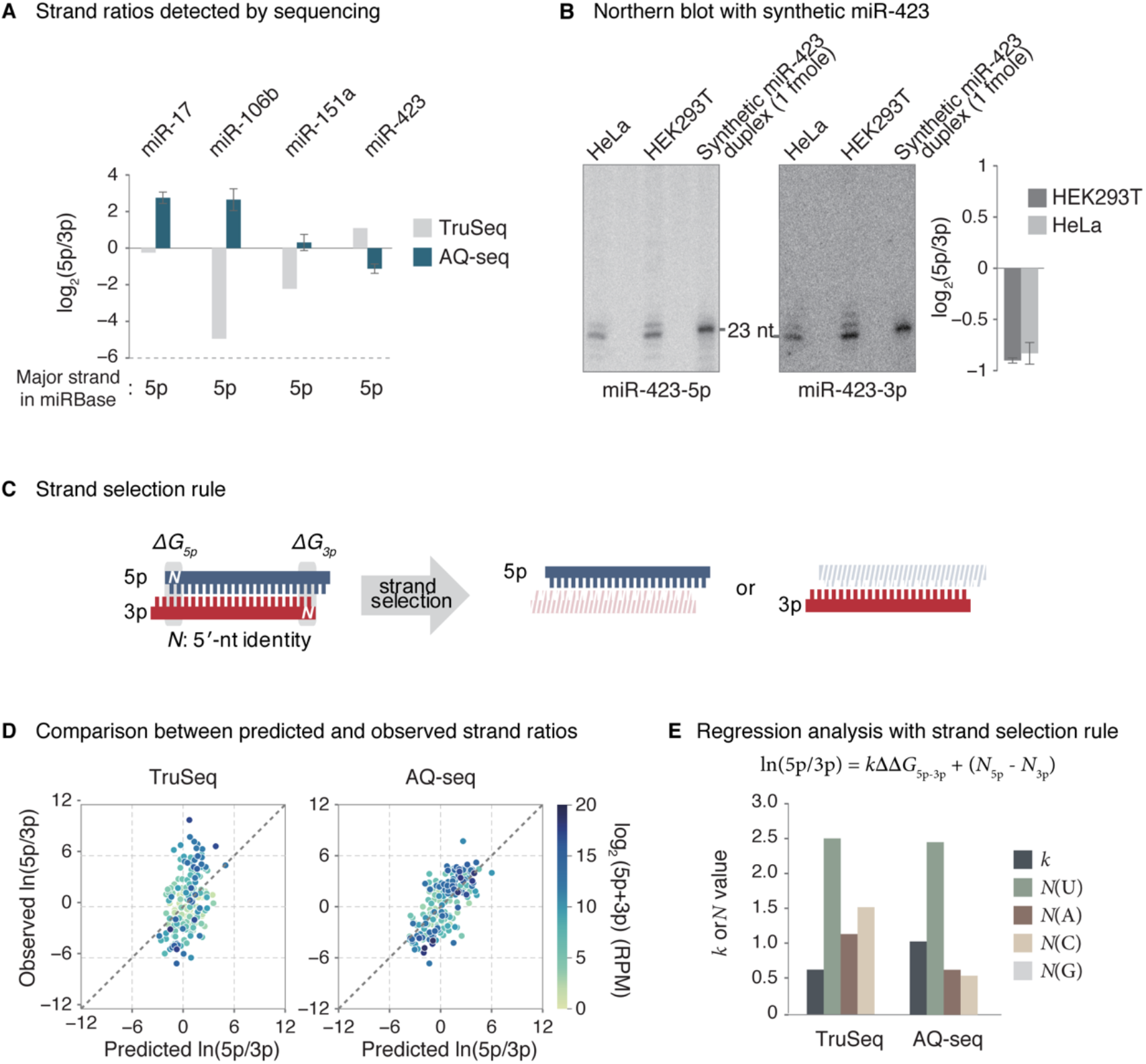
AQ-seq accurately determines strand ratio. (A) Log -transformed 5p/3p ratios obtained from TruSeq and AQ-seq in HEK293T cells. Dominant arms annotated in miRBase are denoted at the bottom. Bars indicate mean +/− standard deviations (s.d.) (n=2). (B) Northern blot of miR-423-5p and miR-423-3p. Left and middle: Total RNAs from HeLa and HEK293T cells were used for miRNA detection. 1 fmol of synthetic miR-423 duplex was loaded for normalization. Right: A bar plot representing log -transformed 5p/3p ratio calculated from band intensities of Northern blot shows that the 3p is the major strand, consistent with AQ-seq data. Bars indicate mean +/− standard deviations (s.d.) (n=2). (C) Two main end properties that determine which strand will be selected: 5′ end identity (N_5p(3p)_) and thermodynamic stability (ΔG_5p(3p)_). (D) Comparison of strand ratios between those predicted by the model and those obtained from TruSeq (left panel) or AQ-seq (right panel) in HEK293T cells. See Methods for details. RPM, reads per million. (E) Linear regression analysis with the indicated equation for strand selection of miRNAs. Bars represent values of the indicated parameters obtained from the model fitted to either TruSeq or AQ-seq data from HEK293T cells.

It has been shown that strand selection is mainly governed by the following rules; (1) The strand whose 5′ end is relatively unstable is favored by AGO and (2) the strand with uridine or adenosine at its 5′ end is selected as a guide strand (Khvorova et al. 2003; Schwarz et al. 2003; Frank et al. 2010; Suzuki et al. 2015). A recent study using systematic biochemical assays reported that strand ratio is predicted well by these rules and can be expressed by the following equation (Fig. 5C):

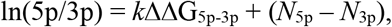

where *k* represents the constant for the relative thermodynamic stability, and *N*_5p_ and *N*_3p_ represent the constants corresponding to the 5′ end nucleotide identity of 5p and 3p strands, respectively (Suzuki et al. 2015).

To assess the performance of AQ-seq in measuring the strand ratio, we fitted the strand selection model to strand ratio values measured by either AQ-seq or TruSeq (Fig. 5D). In the regression analysis, we integrated the 5′ ends defined by AQ-seq for reliable prediction of the strand ratio. We then compared the strand ratios predicted by each fitted model with those experimentally obtained. The predicted strand ratios by the linear model indeed fit better with the AQ-seq data than the TruSeq data (Fig. 5D; Supplemental Fig. S5C). Notably, the fitted model by TruSeq exhibited more dependency on 5′ cytosine and guanosine than thermodynamic stability (Fig. 5E, left; Supplemental Fig. S5D) whereas the model by AQ-seq relies mainly on 5′ uridine and relative thermodynamic stability (Fig. 5E, right; Supplemental Fig. S5D), which conforms with the previously established strand selection rules. Taken together, our analyses indicated that AQ-seq improves the accuracy in measuring arm ratios.

## Discussion

We here establish a protocol, termed AQ-seq, which diminishes the ligation bias of sRNA-seq (Fig. 1A). AQ-seq detects miRNAs of low abundance and reliably defines the terminal sequences of miRNAs undetected when using the conventional sRNA-seq method. Given that AQ-seq data identifies previously undetected termini, it will help us to refine the miRNA annotation in the publicly available databases. It is noteworthy that miRNA target prediction algorithms rely on the complementarity between seed sequences and the targets (Bartel 2009). As the seed sequences are determined by the position relative to the 5′ end of miRNA, incorrect annotation of the 5′ end misleads target identification and functional studies. For instance, the 2-nt shift in the 5′ end of miR-222-5p would result in a distinct set of targets (Fig. 3B). The 1-nt offsets found in miR-192-5p and miR-296-3p are also expected to alter the targets substantially (Fig. 3E). Furthermore, accurate quantification of small RNA will help clean up miRBase because a large fraction of its registry is thought to be false (Wang and Liu 2011; Fromm et al. 2015; Kim et al. 2017). Reliable sRNA-seq data will be imperative to re-evaluate the entries in databases widely used in miRNA research. It is noteworthy that AQ-seq incorporates RNA spike-in controls that consist of 30 exogenous RNA molecules. Use of the spike-ins allows us to monitor ligation bias and detection sensitivity. Therefore, we recommend using spike-ins in sRNA-seq experiments.

The enhanced performance of AQ-seq data gives us an opportunity to build a reliable catalog of isomiRs and thereby improve our understanding of miRNA biogenesis. In this study, we uncover several interesting maturation events. Firstly, we detect alternative processing in 29 miRNAs including miR-101-3p, miR-140-3p, miR-183-5p, miR-192-5p, and miR-296-3p. By generating multiple isomiRs from a single hairpin, miRNAs can expand their regulatory capacity. It will be interesting in future studies to investigate whether or not alternative processing of miRNA is regulated in a condition-specific manner, as in the cases of pre-mRNA splicing and alternative polyadenylation. Secondly, we find that more than 15 miRNAs are uridylated at a frequency of over 30% in HEK293T or HeLa cells, and that TUT4/7 mediate the uridylation at the pre-miRNA stage. This indicates that uridylation influences the miRNA pathway beyond the let-7 family. It will be of interest to study the functional consequences of uridylation of these miRNAs. Lastly, we accurately measure the strand ratio, which will be critical for the studies of alternative strand selection or “arm switching” whose mechanism remains unknown.

Growing evidence suggests that isomiRs are generated in a context-dependent manner and play physiologically relevant roles during developmental and pathological processes (Hinton et al. 2014; Tan et al. 2014; Guo et al. 2016; Wang et al. 2016). Notably, it was shown that isomiR profiles successfully classify diverse cancer types and, when combined with miRNA profiles, improve discrimination between cancer and normal tissues (Telonis et al. 2015; Koppers-Lalic et al. 2016; Telonis et al. 2017), highlighting isomiRs as potential diagnostic biomarkers. Therefore, applying AQ-seq to biosamples may enhance the diagnostic power of sRNA-seq.

It is also important to note that AQ-seq can be adopted to other library preparation methods which involve adapter ligation. In particular, AQ-seq would be beneficial to studies which require “within-sample” comparisons. For example, crosslinking and immunoprecipitation followed by sequencing (CLIP-seq) identifies the protein-RNA interaction sites by quantitatively detecting crosslink sites (Konig et al. 2012). Since the analysis takes into account relative enrichment of crosslink sites, it is crucial to quantify and compare the abundance of the individual sites within a given sample, which can be severely compromised by ligation bias. Therefore, the inclusion of randomized adapters and PEG in CLIP-seq experiments will significantly improve the identification of true binding sites. Another potentially useful application is with ribosome profiling (Ribo-seq) which detects ribosome-protected mRNA fragments (RPFs) by sequencing (Ingolia et al. 2009). Ribo-seq is used widely to systematically investigate the mechanism and regulation of translation and the identification of open reading frames. This technique requires unbiased capture of RPFs, which may be achieved by applying randomized adapters and PEG as shown in this study. Taken together, we anticipate AQ-seq will serve as a powerful tool that contributes not only to small RNA studies but also to RNA researches that depend on ligation-based library generation.

## Methods

### Small RNA sequencing library preparation

Total RNA was isolated from HEK293T and HeLa cells using TRIzol (Invitrogen) and mixed with 1 μL of 10 nM spike-in control oligos, 30 non-human RNA sequences of 21-23 nt in length (Supplemental Table S1). The oligos were synthesized from Bioneer Inc., resuspended in distilled water and pooled at equimolar concentrations. The RNA mixture was size-fractionated by 15% urea-polyacrylamide gel electrophoresis (PAGE) to enrich miRNA species using two FAM-labeled markers (17 nt and 29 nt). Small RNA libraries were constructed using either TruSeq small RNA library preparation kit (Illumina) according to manufacturer’s instructions or AQ-seq methods as follows: miRNA-enriched RNA was ligated to the 3′ randomized adapter using T4 RNA ligase 2 truncated KQ (NEB) in the presence of 20% PEG 8000 (NEB). The ligated RNA was gel-purified on a 15% urea-polyacrylamide gel using two markers (40 nt and 55 nt) to remove the free 3′ adapter. The purified RNA was ligated to the 5′ randomized adapter using T4 RNA ligase 1 (NEB) in the presence of 20% PEG 8000 (NEB). The products were reverse-transcribed with RT primer (RTP, TruSeq kit; Illumina) using SuperScript III reverse transcriptase (Invitrogen) and amplified with primers (RP1 forward primer and RPIX reverse primer, TruSeq kit; Illumina) using Phusion High-Fidelity DNA Polymerase (Thermo Scientific). The PCR-amplified cDNA was gel-purified to remove adapter dimers and sequenced using MiSeq or HiSeq platforms. Markers for size-fractionation and randomized adapters were synthesized from IDT and are listed in Supplemental Table S2.

### Analysis of small RNA sequencing

The TruSeq 3′ adapter sequence was removed from FASTQ files using cutadapt (Martin 2011). For AQ-seq data, 4 nt-long degenerate sequences at the 3′ and 5′ end were trimmed using the FASTX-Toolkit (http://hannonlab.cshl.edu/fastx_toolkit/). Next, reads shorter than 18 nt were filtered out, and then low-quality reads (phred quality <20 or <30 in >95% or >50% of nucleotides) or artifact reads were discarded with the FASTX-Toolkit.

Preprocessed reads were first aligned to the spike-in sequences by BWA with an option of –n 3 (Li and Durbin 2010). Reads mapped to spike-in sequences perfectly or with a single mismatch were considered as reliable spike-in reads. Proportions of read counts for each spike-in were used for the representation of sRNA-seq bias.

Reads unmapped to spike-in sequences by BWA with an option of –n 3 were subsequently mapped to the human genome (hg38) with the same option. For multi-mapped reads, we selected the alignment results which have the best alignment score, allowing mismatches only at the 3′ end of reads using custom scripts. Reads overlapped with the genomic coordinates of miRNAs (from miRBase v21) were identified by intersectBed in BEDTools and used for further analysis (Quinlan and Hall 2010; Kozomara and Griffiths-Jones 2014).

### Primer extension

The primer was labeled at the 5′ end with T4 polynucleotide (Takara) and [γ-^32^P] ATP. RNA was extracted using TRIzol reagent (Invitrogen) and then small RNA fraction was enriched using mirVana kit (Ambion). The RNA samples were reverse-transcribed with a 5′ end-radiolabeled primer using SuperScript III reverse transcriptase (Invitrogen). The products were separated on a 15% urea-polyacrylamide gel and the radioactive signals were analyzed using a BAS-2500 (FujiFilm). The sequence of RT primer is listed in Supplemental Table S2.

### Northern blot analysis

RNA was isolated using either TRIzol (Invitrogen) or mirVana kit (Ambion), resolved on a 15% urea-polyacrylamide gel, transferred to a Hybond-NX membrane (Amersham) and then crosslinked to the membrane chemically with 1-ethyl-3-(3-dimethylaminopropyl) carbodiimide (Pall and Hamilton 2008). The probes were labeled at the 5′ end with T4 polynucleotide (Takara) and [γ-^32^P] ATP. 5′ end-radiolabeled oligonucleotide complementary to the indicated miRNA was hybridized to the membrane. The radioactive signals were analyzed using a BAS-2500. Band intensities were quantified by Multi Gauge software. To strip off probes, the blot was incubated with a pre-boiled solution of 0.5% SDS for 15 min. Synthetic miRNA duplexes were synthesized by Bioneer (AccuTarget). The sequences of probes are listed in Supplemental Table S2.

### Quantitative real-time PCR

Total RNA was reverse-transcribed using TaqMan miRNA Reverse Transcription kit (Applied Biosystems), and subjected to quantitative real-time PCR with TaqMan gene expression assay kits (Applied Biosystems) according to manufacturer’s instructions. U6 snRNA was used for internal control.

### Prediction of miRNA arm ratio

We analyzed miRNA arm ratio as described in (Suzuki et al. 2015) with the following model (Fig. 5C):

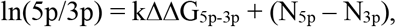

where k and N_5p(3p)_ represent the constant for the relative thermodynamic stability and the constant corresponding to the 5′ end identity, respectively.

For prediction, we constructed miRNA duplexes using sequences defined by AQ-seq. If two replicates generated by AQ-seq method reported the identical ends, and the end identities were inconsistent with miRBase, we replaced the miRBase sequences with the those defined by AQ-seq. RNA secondary structure was folded by mfold version 3.6 and thermodynamic stability (ΔG_5p(3p)_) with dinucleotide subsequences was estimated based on the mfold result (Zuker 2003). Next, we selected top 200 abundant miRNAs 1) which are not duplicated in the genome, 2) whose 5p and 3p are annotated in miRBase, and 3) whose mature sequences had homogenous 5′ ends (miRNA duplexes whose predominant 5′ ends from 5p and 3p strands accounted for more than 90% of total reads (5p +3p)). We measured the log ratios of 5p/3p strands using either TruSeq or AQ-seq and performed regression analysis with the previously described equation in R.

## Data access

The small RNA sequencing data in this study will be uploaded to a publicly available database such as GEO before publication.

## Acknowledgements

We are grateful to Eunji Kim for technical help. We thank Jaechul Lim, Boseon Kim, Young Yoon Lee, Young-suk Lee, Hyunjoon Kim, Yongwoo Na, and other members of our laboratory for discussion. This work was supported by IBS-R008-D1 from the Institute for Basic Science from the Ministry of Science, ICT and Future Planning of Korea (H.K., J.K., K.K., H.C., K.Y., and V.N.K.), by the BK21 Research Fellowships from the Ministry of Education of Korea (H.K. and K.K.), and by NRF (National Research Foundation of Korea) Grant funded by Korean Government (NRF-2015-Global Ph.D. Fellowship Program) (H.K.).

## Author contributions

H.K., J.K., K.K., K.Y., and V.N.K. designed experiments. H.C. designed small RNA spike-ins. J.K. and K.Y. generated sRNA-seq libraries. H.K. and J.K. performed biochemical experiments. H.K. carried out computational analyses. H.K., J.K., and V.N.K. wrote the manuscript.

